# Anti-Kv1.2 Immunoprecipitation Identifies shared mGluR1-Associated Signalosome Complex proteins and PKC-Mediated Regulation of Kv1.2 in the Cerebellum

**DOI:** 10.1101/2025.11.09.687499

**Authors:** Sharath C. Madasu, Anthony D. Morielli

## Abstract

The voltage gated potassium channel Kv1.2 plays a key role in the central nervous system and mutations in Kv1.2 leads to neurological disorders such as epilepsies and ataxias. In the cerebellum regulation of Kv1.2 is coupled to learning and memory. We have previously shown that blocking Kv1.2 by infusing its specific inhibitor Tityustoxin-kα (TsTX) into the lobulus simplex of the cerebellum facilitates eyeblink conditioning (EBC) and that EBC modulates Kv1.2 surface expression in cerebellar interneurons. The metabotropic glutamate receptor mGluR1 is required for EBC although the molecular mechanisms are not fully understood. We have previously shown that infusion of the mGluR1 agonist (S)-3,5-dihydroxyphenylglycine (DHPG) into the lobulus simplex of the cerebellum mimics the faciliatory effect of TsTX on EBC. We therefore hypothesize that mGluR1 could act, in part, through suppression of Kv1.2. Earlier studies have shown that Kv1.2 suppression involves channel tyrosine phosphorylation and its endocytocytic removal from the cell surface. In this study we report that an excitatory chemical stimulus (50mM K^+^-100µM glutamate) applied to cerebellar slices enhanced Kv1.2 tyrosine phosphorylation and that this increase was lessened in the presence of the mGluR1 inhibitor YM298198. More direct evidence for mGluR1 modulation of Kv1.2 comes from our finding that selective activation of mGluR1 with DHPG reduced the amount of Kv1.2 detected by cell surface biotinylation in cerebellar slices. To determine the molecular pathways involved we used an unbiased mass spectrometry-based proteomics approach to identify Kv1.2-protein interactions that are modulated by mGluR1. Among the interactions enhanced by DHPG were those with PKC-γ, CaMKII and Gq/G11, each of which had been shown in other studies to co-immunoprecipitate with mGluR1 and contribute to its signaling. Of particular note was the interaction between Kv1.2 and PKC-γ since in HEK cells and hippocampal neurons Kv1.2 endocytosis is elicited by PKC activation. Here we show that activation of PKCs with PMA reduced surface Kv1.2, while the PKC inhibitor Go6983 attenuated the reduction in surface Kv1.2 levels elicited by DHPG, suggesting that the mechanism by which mGluR1 modulates cerebellar Kv1.2 likely involves PKC.

## Introduction

Purkinje cells (PCs) are the sole output of the cerebellar cortex and a major site of signal integration for motor coordination and motor learning. Khavandagar *et al* showed that dendrotoxin-sensitive voltage gated potassium channel (Kv1) play an important role in regulating the excitability of PCs by preventing the occurrence of random calcium spikes and hyperexcitability of PCs. Kv1.2 is highly expressed in PCs (1,2) and regulates their excitability. Basket cells (BC) are inhibitory interneurons in cerebellar cortex that synapse with PCs and Kv1.2 is also expressed in the basket cell axon terminal where it regulates the release of the inhibitory neurotransmitter GABA. Therefore, regulation of Kv1.2 ion channels in the PC and BC influences the excitability of PC and therefore the output of the cerebellum.

Genetic mutation studies have shown that I402T missense mutation of Kv1.2 in the cerebellum decreases Kv1.2 surface expression in the basket cell axon terminal and increases firing of basket cells, reducing the firing frequency of PCs. These mouse models showed movement disorders including abnormal gait, measured in coordinated footsteps, that indicate cerebellar deficiencies (3). Kv1.2 knockout animal models show seizures and reduced life span (4) and mutations in Kv1.2 have been found to cause severe deficits in growth, development, ataxia, epilepsies in mouse models and seizures in human patients (5–8). These findings suggest that regulation of Kv1.2 could play an important physiological role in shaping neuronal excitability and overall health.

An important mechanism of Kv1.2 regulation involves endocytic trafficking to and from cell surface, we have previously shown that the surface expression of Kv1.2 is regulated by tyrosine and serine/threonine kinases (2,9–12). In the cerebellum, Kv1.2 expression at the cell surface is regulated by the secretin receptor through a process involving protein kinase A (PKA)(13). Such regulation likely contributes to a range of cerebellar functions, including learning. Infusion of secretin into the lobulus simplex region of the cerebellar cortex enhanced eyeblink conditioning, and this effect was mimicked by infusion of the specific Kv1.2 inhibitory peptide Tityustoxin-kα (TsTX)(13). A later study showed that EBC alters the surface expression of Kv1.2 in regions of the cerebellar cortex (14). Together, these studies indicate that regulation of Kv1.2 expression at the cell surface is a component of the complex mechanisms underlaying cerebellum-dependent learning.

mGluR1, a G-protein-coupled receptor which is highly expressed in PCs is required for EBC. Knock out of mGluR1 impairs EBC (15–19), mGluR1 rescue restored EBC in these knockout mice (16,18). Moreover, mGluR1 knockout mice also exhibited impaired parallel fiber – PC cell long term depression, indicating that mGluR1 is important for PC synaptic plasticity (16).

Given that both mGluR1 and Kv1.2 affect PC excitability and Kv1.2 regulation is associated with EBC, we hypothesized that mGluR1 may regulate Kv1.2 in the cerebellum. We previously have shown that mGluR1 activation with the specific mGluR1/5 agonist (S)-3,5-Dihydroxyphenylglycine (DHPG) enhances EBC in adult rats. An LTD inducing stimulus which known to activate mGluR1, increases tyrosine phosphorylation on Kv1.2 and some of its co-immunoprecipitated proteins. Using a cell surface biotinylation assay we show that mGluR1 activation via DHPG and PKC activation via PMA reduces surface Kv1.2 in cerebellar slices. To identify molecular pathways involved in this regulation we took an unbiased mass spectrometry approach and found that known interactors of mGluR1 such as PKC-γ, calmodulin kinase (CaMKII) and Gq/11 (20) co-immunoprecipitate with Kv1.2 and that those interactions are altered by mGluR1 activity. Using pharmacological inhibitor of PKC Go6983 (PKCi in figures) we observed that PKC inhibition attenuates decrease in surface Kv1.2 mediated by mGluR1 and PMA, suggesting that mGluR1 affects Kv1.2 at least in part through a PKC-dependent mechanism.

## Materials & Methods

### Antibodies

Kv1.2 antibodies clone K14/16 from Neuromab UC Davis. While Pan-CaMKII, AMPAR2, p-Tyr 100, PKA p-Ser/Thr substrate antibody and Calbindin were purchased from Cell signaling. PKC - γ and p-PKC - γ thr514 antibodies were purchased from Genetex, Grid2 extracellular Antibody was from Alomone labs. Anti – mouse Dx700 was purchased from Invitrogen and anti – rabbit 800 infrared dye conjugated 2° antibodies were purchased from Rockland INC.

### Chemicals

DHPG, Go6983, EGLU, NCQX, DAP –V, tetrodotoxin were purchased from Tocris and all other chemicals were purchased from sigma unless specified. HEPES was purchased from Affymetrix, Sulfo NHHS-S-S-Biotin, HALT protease inhibitor, Neutravidin beads were purchased from Thermofisher. BPV-phen, mg-132 were purchased from Millipore.

### Slice preparation

400 μm thick cerebellar slices were generated from cerebellum of 4-6-week-old male Sprague dawley rats using a Lecia 1200 vibratome using an adaptation of the protective recovery method (21). Briefly, slices were generated in ice-cold Holding and Slicing ACSF, allowed to recover in Recovery ACSF at 34°C for 7 min and then transferred to Holding and Slicing ACSF at 25°C for at least 1 hour. **Recovery ACS**F: (quantity in milli moles mM), 93 NMDG (N-Methyl D Glucosamine), 2.5 KCl, 1.2 NaH_2_PO_4_, 30 NaHCO_3_, 20 HEPES, 25 glucose, 5 sodium ascorbate, 2 thiourea, 3 sodium pyruvate, 12 N-acetyl cysteine, pH adjusted to 7.4 then 10 MgSO_4_*7H_2_O, 0.5 CaCl_2_. **Holding and Slicing ACSF**: 92 NaCl, 2.5 KCl, 1.2 NaH_2_PO_4_, 30 NaHCO_3_, 20 HEPES, 25 glucose, 5 sodium ascorbate, 2 thiourea, 3 sodium pyruvate, 12 N-acetyl cysteine, pH adjusted to 7.4 then 10 MgSo_4_.7H_2_O, 0.5 CaCl_2_. **Recording ACSF**: 124 NaCl, 2.5 KCl, 1.2 NaH_2_PO_4_, 24 NaHCO_3_, 5 HEPES, 12.5 glucose, 2 MgSo_4_.7H_2_O, 2 CaCl_2_. pH 7.4. The osmolarity of all ACSF solutions was adjusted to 290 milliosmoles.

### Drug treatment

The slices were then treated with 50 μM DHPG for 10 min or 1 μM PMA 30 min in presence of 100nM TTX, (in µM) 100 EGLU, 50 DAP-V, 100 NCQX made recording ACSF and were incubated for 1 hour in ACSF minus DHPG after drug treatment. For PKC inhibition, the slices were pre-incubated with 1μM PKC inhibitor Go6983 (PKCi in figures) for 1 hour then the respective DHPG (10 min) or PMA (30 min) was added in presence of Go6983. After 10 min, the DHPG was replaced with recording ACSF containing the Go6983. After the drug treatment, the slices were put in Ice cold HBSS to stop endocytosis and were biotinylated as below.

### Biotinylation

2mg/ml Sulfo-NHS-SS-Biotin was made up in ice-cold HBSS (thermos) and once slices were treated with drugs for indicated times, biotin was applied to slices for 15 min (all in Ice). Then the unreacted biotin was quenched using ice cold 50mM tris buffered HBSS (pH 7.5). The slices were then transferred to RIPA (containing protease inhibitor HALT 1X, 1mM DTT, 10µM proteasome inhibitor Mg132, Sodium Fluoride 100µM, 2mM EDTA, Sodium Orthovanadate 100µM, 5mM NEM, 0.1µM BpV-Phen) and were lysed by sonication using brief 8 sec pulses twice. The 800 µL of lysate was incubated with 40μl of nutravidin bead slurry overnight to separate surface proteins, the remaining 200µL lysate was saved as total protein. The nutravidin bound proteins were washed 3 times with RIPA for 10 min and were eluted by boiling in lamelli buffer containing 100 mM DTT and were subjected to electrophoresis 10% Bis-poly acrylamide gel. Then total and surface levels of Kv1.2, AMPAR2 (1:1000) and calbindin (1:2000) were probed by western blots. To estimate surface protein a ratio of the integrated intensities (Li-Cor Odyssey) of biotinylated Kv1.2 to total Kv1.2 was measured. The total Kv1.2 was obtained by taking the ratio of integrated intensities of total Kv1.2 (of individual slices) to the average of total Kv1.2 across the blot to account for blot to blot variability. The surface protein ratio from individual experiments were normalized to control. So, the controls are averaged to 1 indicating 100% surface channel. All such individual experiments were pooled and averaged together, so N represents individual experiments and n represents number of slices in each condition.

### Inclusion criteria

Calbindin was used as intracellular marker. Only the slices which had no calbindin signal in the eluted or surface portion were considered healthy non permeabilized slices and were used for quantification.

### Western Blots

Proteins were transferred on to nitro-cellulose membrane and were blocked by 5% BSA and strips separating molecular weights of 55KD-90KD to be probed using mouse anti - Kv1.2(2µg), rabbit anti - AMPAR2 (1:1000) and 10KD – 55KD to be probed using rabbit anti – Calbindin (1:2000) antibodies overnight. The membranes were then washed with TBST and incubated with anti – mouse Dx700 and anti – rabbit 800 infrared dye conjugated 2° antibodies. The images were developed using Li-cor odyssey infrared scanner in its linear range of detection. The blots were quantified using the accompanying Odyssey software and integrated intensities of individual bands were measured for quantification. Calbindin was used as a cytosolic protein marker indicating live and healthy slices.

### Membrane fractionation and anti Kv1.2 immunoprecipitation

Slices were generated and recovered as described above. The slices were then treated with 100 µM DHPG or PBS for 10 min or 60 min and were snap frozen in liquid nitrogen and stored in −80°C until further use slices form 6 - 8 different animals were pooled in an experiment according to their treatment conditions. The frozen slices were then homogenized using a Dounce homogenizer in ice cold HEPES buffer at 1ml/100mg of brain tissue containing HALT 1X, 1mM DTT, 10µM proteasome inhibitor Mg132, 100µM Sodium Fluoride, 2mM EDTA, 100µM Sodium Orthovanadate, 5mM NEM, 0.1µM BpV-Phen, 5 mM BATA. The homogenates were subjected to centrifugation at 1000g for 15 min to remove the nuclear pellet. The supernatant was further subjected to centrifugation at 200,000g for 30 min at 4°C and resulting pellet was resuspended in half of the initial volume of HEPES buffer and was re-centrifuged. At the end of second centrifugation the pellet was resuspended in 1 ml lysis buffer and was centrifuged at 1000g’s for 10 min to remove insoluble contents. The supernatant contains the membrane fraction and is used for immunoprecipitation. 80µg of Kv1.2 antibodies were cross linked to 8mg of epoxy coated Dyna magnetic beads according to manufacturer’s instructions. Equal amount of protein from the membrane fraction was incubated with the crosslinked antibody bead complex overnight (10 – 16 hours). The immunoprecipitated proteins were separated washed thrice for 10 min at 4°C with lysis buffer. The proteins were eluted using high pH buffer as per manufacturer’s instructions and the eluate boiled in Lammeli buffer with 100mM DTT and were separated on 10% SDS poly acrylamide gel and stained with Coomassie Blue.

### Mass Spectrometry

#### Trypsin digestion and peptide labeling by tandem mass tag (TMT)

The gel regions containing the separated proteins but not the antibody heavy and light chains for each condition were minced, combined and subjected to disulfide reduction using 10mM DTT in 100mM triethylammonium bicarbonate (TEAB, Thermo) for 1 hour at 56^0^C. The reduced cysteines were then alkylated with 55mM iodoacetamide for 45 min at room temperature in the dark. The gel pieces were washed with TEAB for 10 min at room temperature, dehydrated with 100% acetonitrile and washed with TEAB for 10 min and were dehydrated again in 100% acetonitrile followed by in-gel trypsin digestion (6 ng/µl of sequencing grade trypsin enough to cover entire gel piece) overnight at 37°C. The digested peptides were extracted by incubation in 5% formic acid for 1 hour, followed by incubation in 5% formic acid in 50% acetonitrile. The extracted peptides were subjected to evaporation in a SpeedVac. The peptides from each condition were labeled with one of the isobaric tags from the TMT-sixplex (Thermo Fisher) according to the manufacturer’s instructions. Briefly, the peptides resuspended in 102.5 uL TEAB were incubated with 41 uL TMT reagents for 1 h at room temperature (see the table insert in Figure 3a for the information on specific tags that were used) after which 8 μl of 50% hydroxylamine was added to quench the reaction. Equal amount of the labelled peptides were pooled and dried down. The combined TMT-labeled peptides were resuspended in 10 μl of 2.5% acetonitrile and 2.5% formic acid.

#### Liquid chromatography-tandem mass spectrometry (LC-MS/MS)

The purified labeled and combined peptides were resuspended in 2.5% acetonitrile (CH3CN) and 2.5% formic acid (FA) in water for subsequent LC-MS/MS based peptide identification and quantification. Analyses were performed on the Q-Exactive mass spectrometer coupled to an EASY-nLC (Thermo Fisher Scientific, Waltham, MA, USA). Samples were loaded onto a 100 μm x 120 mm capillary column packed with Halo C18 (2.7 μm particle size, 90 nm pore size, Michrom Bioresources, CA, USA) at a flow rate of 300 nl min-1. The column end was laser pulled to a ∼3 μm orifice and packed with minimal amounts of 5um Magic C18AQ before packing the column with the 3-μm particle size chromatographic materials. Peptides were separated using a gradient of 2.5-35% CH3CN/0.1% FA over 150 min, 35-100% CH3CN/0.1% FA in 1 min and then 100% CH3CN/0.1% FA for 8 min, followed by an immediate return to 2.5% CH3CN/0.1% FA and a hold at 2.5% CH3CN/0.1% FA. Peptides were introduced into the mass spectrometer via a nanospray ionization source and a laser pulled ∼3 μm orifice with a spray voltage of 2.0 kV. Mass spectrometry data was acquired in a data-dependent “Top 10” acquisition mode with lock mass function activated (*m/z* 371.1012; use lock masses: best; lock mass injection: full MS), in which a survey scan from *m/z* 350-1600 at 70, 000 resolution (AGC target 1e6; max IT 100 ms; profile mode) was followed by 10 higher-energy collisional dissociation (HCD) tandem mass spectrometry (MS/MS) scans on the most abundant ions at 35,000 resolution (AGC target 1e5; max IT 100 ms; profile mode). MS/MS scans were acquired with an isolation width of 1.2 *m/z* and a normalized collisional energy of 35%. Dynamic exclusion was enabled (peptide match: preferred; exclude isotopes: on; underfill ratio: 1%; exclusion duration: 30 sec).

#### Data analysis

Product ion spectra were searched using SEQUEST and Mascot implemented on the Proteome Discoverer 2.2 (Thermo Fisher Scientific, Waltham, MA, USA) in the processing workflow against a curated Uniprot Rattus norwegicus protein database (AUP000000537; downloaded on June 9, 2017). Search Parameters were as follows: (1) full trypsin enzymatic activity; (2) mass tolerance at 10 ppm and 0.02 Da for precursor ions and fragment ions, respectively; (3) dynamic modifications: oxidation on methionine (+15.995 Da), and phosphorylation on serine, threonine, and tyrosine (+79.96633 Da); (4) dynamic TMT6plex modification on N-termini and lysine (+229.163 Da); and (5) static carbamidomethylation modification on cysteines (+57.021 Da). Percolator node was included in the workflow to limit the false positive (FP) rates to less than 1% in the data set. The relative abundances of TMT labeled peptides were quantified with the Reporter Ions Quantifier node in the consensus workflow and parameters were set as follows: (1) both unique and razor peptides were used for quantification; (2) Reject Quan Results with Missing Channels: False; (3) Apply Quan Value Corrections: False; (4) Co-Isolation Threshold: 50; (5) Average Reporter S/N Threshold = 10; (6) “Total Peptide Amount” was used for normalization and (7) Scaling Mode was set “on All Average”. The ProteinCenter Annotation workflow node was included for Biological Process/Cellular Component/Molecular Function and Wiki/KEGG Pathway annotations. All the protein identification and quantification information (<1% FP; with protein grouping enabled) was exported from the “.msf” result files to Excel spreadsheets (Supplementary Table 1) for further statistical analyses.

Proteins that were identified in both experiments were kept and all protein abundance ratios (DHPG/Veh) in each condition were normalized to the corresponding Kv1.2 ratio. Average fold change (DHPG/Veh at 1h vs. 10 min) was calculated across the two independent biological replicates with two technical replicates and Student’s t-test was performed (Supplementary Table). For proteins with a p value < 0.05, the normalized ratios were imported into the JMP Pro 13 (SAS Institute, Cary, North Carolina, US) to construct the heatmaps.

### Immunofluorescence

40µM slices were obtained from 3-4-week-old rats which were perfused with2% formaldehyde in PBS. The slices were then incubated in 0.3% triton in NGS PBS (at Rt) for epitope unmasking. The slices were then washed and blocked in 3% NGS in PBS solution for 1 hour followed by overnight incubation at room temp with primary antibodies at these dilutions: rabbit anti - mGluR1 (1:50), rabbit anti-PKC γ (thr514) (1:50), rabbit anti calbindin (1:50) mouse anti - Kv1.2 (1:200) in 3% NGS PBS in dark at Rt. The slices were washed in ice cold PBS thrice and were incubated with secondary antibodies goat anti mouse Alexa 647 (1:500), Alexa anti rabbit 568(1:500) overnight and 300nM DAPI nuclear stain for 5 min. The slices were then washed thrice and mounted using prolong Gold antifade solution. The slices were then observed on a C-2 confocal microscope from Nikon instruments couple to an Olympus inverted microscope with settings to collect images in the z-stack at a step size of 0.856, pixel dwell time setting at 5.5, the images were captured using 40x oil immersion objective at a resolution of 2048 x 2048. The laser strength and HV setting were adjusted individually for each channel. The 568 and DAPI images were collected together followed by 647 for the same slice with the same Z co-ordinates. The images were then processed using Fiji image editor. The Z-stacks were compressed and merged to generate the composite image.

## Results

### mGluR1 activation and increases Tyrosine phosphorylation on Kv1.2

Keiko Tanaka *et al* and Augustine *et al* have shown that a chemical LTD process involving depolarization with potassium and glutamate initiates the LTD cascade by activating mGluR1 in cerebellar slices(24). They identify a brief excitatory period of PC excitation lasting 5-10 minutes, followed by LTD initiation after 20-30 minutes (24). Chae *et al* (25) have showed that such a stimulus also results in endocytosis of AMPAR2, via phosphorylation of S880 (26,27). Surface expression of Kv1.2 is also regulated by phosphorylation dependent mechanisms (2,12,14,23,28,29), involving serine/threonine and tyrosine kinases (10,11,13). Since Kv1.2 surface expression both influences and is influenced by EBC, we predicted that chemical LTD stimulus would modulate Kv1.2 phosphorylation levels. Cerebellar slices were treated with a K-Glu stimulus for 5 min (50 mM potassium and 100µM glutamate). Picrotoxin (100µM) was included to reduce inhibitory drive during the stimulus. A subset of slices were pre-treated with the mGluR1 inhibitor YM298198(YM). Slices were collected after 5 minutes and processed for immunoprecipitation and immunoblot as described in Methods. Blots of immunoprecipitated Kv1.2 were probed with anti-phosphotyrosine and anti-PKA substrate antibodies. In the pre-immunoprecipitation lysate we observed an increased ser/thr phosphorylation of proteins (100 – 130 Kd range) with K-Glu treatment for 5 min (normalized intensity K-Glu = 2.27± 0.298 vs Vehicle, N=3, **p= 0.0081 Dunnett’s multiple comparison) indicating that the slices actively respond to the stimulus. Inhibition of the response by the mGluR1 inhibitor YM (Figure 2A) indicates a role for mGluR1 in this process. Using p-Tyrosine 100 antibody we observed and increase in tyrosine phosphorylated proteins, with the strongest signal in the molecular weight range expected for Kv1.2 (Figure 2B) (Relative intensity K-Glu = 1.507 vs Vehicle, *p = 0.021, students T test, intensity was measured from 55 Kd to around 170KD). mGluR1 inhibitor YM attenuates such increase in binding of tyrosine phosphorylated proteins to Kv1.2.

**Figure 1:**
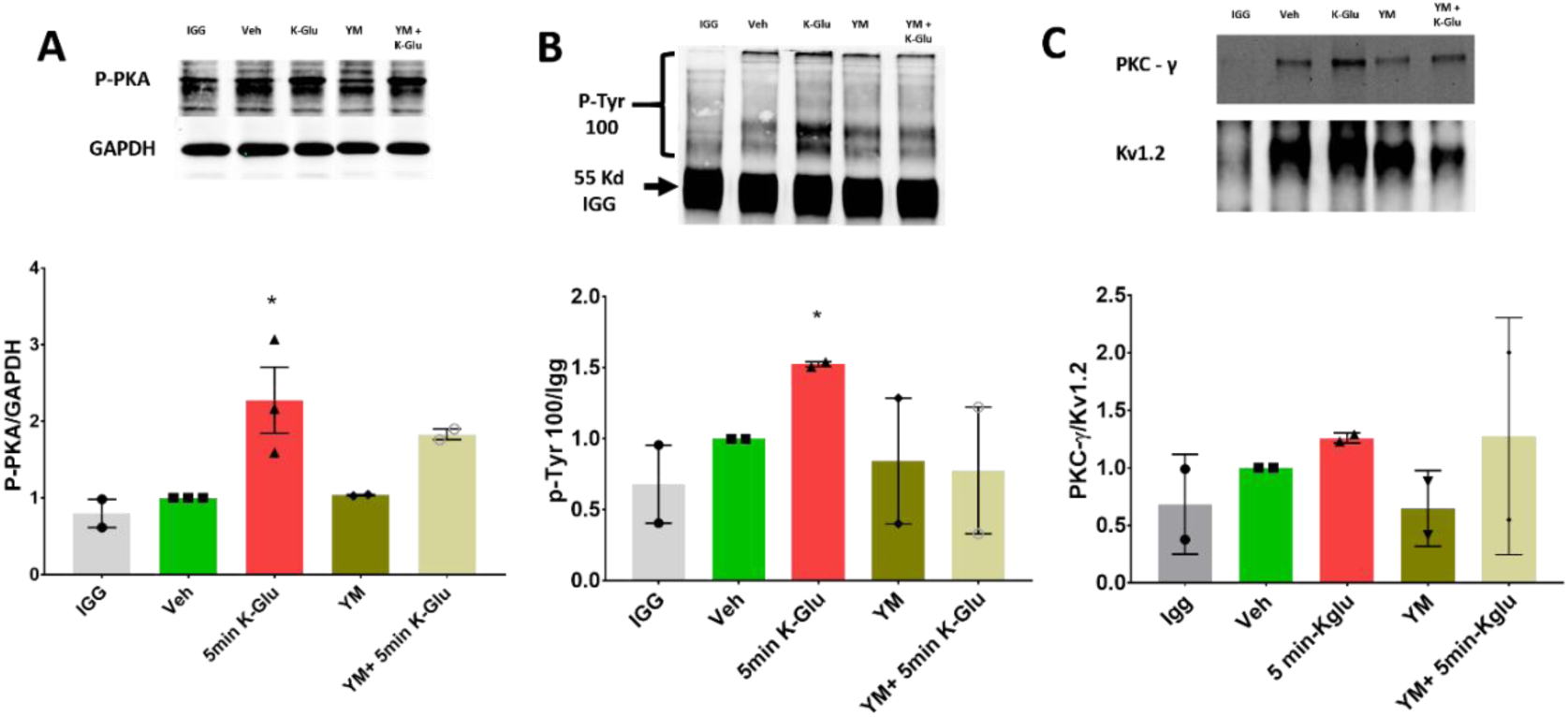
Identification of Kv1.2 Tyrosine Phosphorylation: Cerebellar slices were treated with 50mM Potassium+100 µM Glutamate (K-Glu) or vehicle, in absence or presence of mGluR1 inhibitor YM298198 for 5 min and slices were collected after 1-hour post treatment or after 5 min of K-Glu treatment and were subjected to western analysis with anti – PKA ser/thr substrate antibody. **A)** K-Glu treatment increased Ser/Thr phosphorylation of proteins around 130 KD with in 5 min of treatment even in presence of YM298198. below bar graph quantifying the intensities relative to Vehicle treated slices N=3, **p= 0.0081 vs Vehicle Dunnet’s multiple comparison test. **B)** Cerebellar slices treated with K-Glu were subjected to Kv1.2 immunoprecipitation and were subjected to western analysis using anti p-Tyr antibody, K-Glu treatment results in increase in tyrosine phosphorylation (bar graph below), YM298198 attenuates such increase. N=2, *p=0.021 vs Vehicle student’s T test, Dunnett’s multiple comparison test p = 0.857 vs Vehicle. **C)** Western blot analysis of the IP was probed for PKC-γ (top) and Kv1.2 (below).

**Figure 2:**
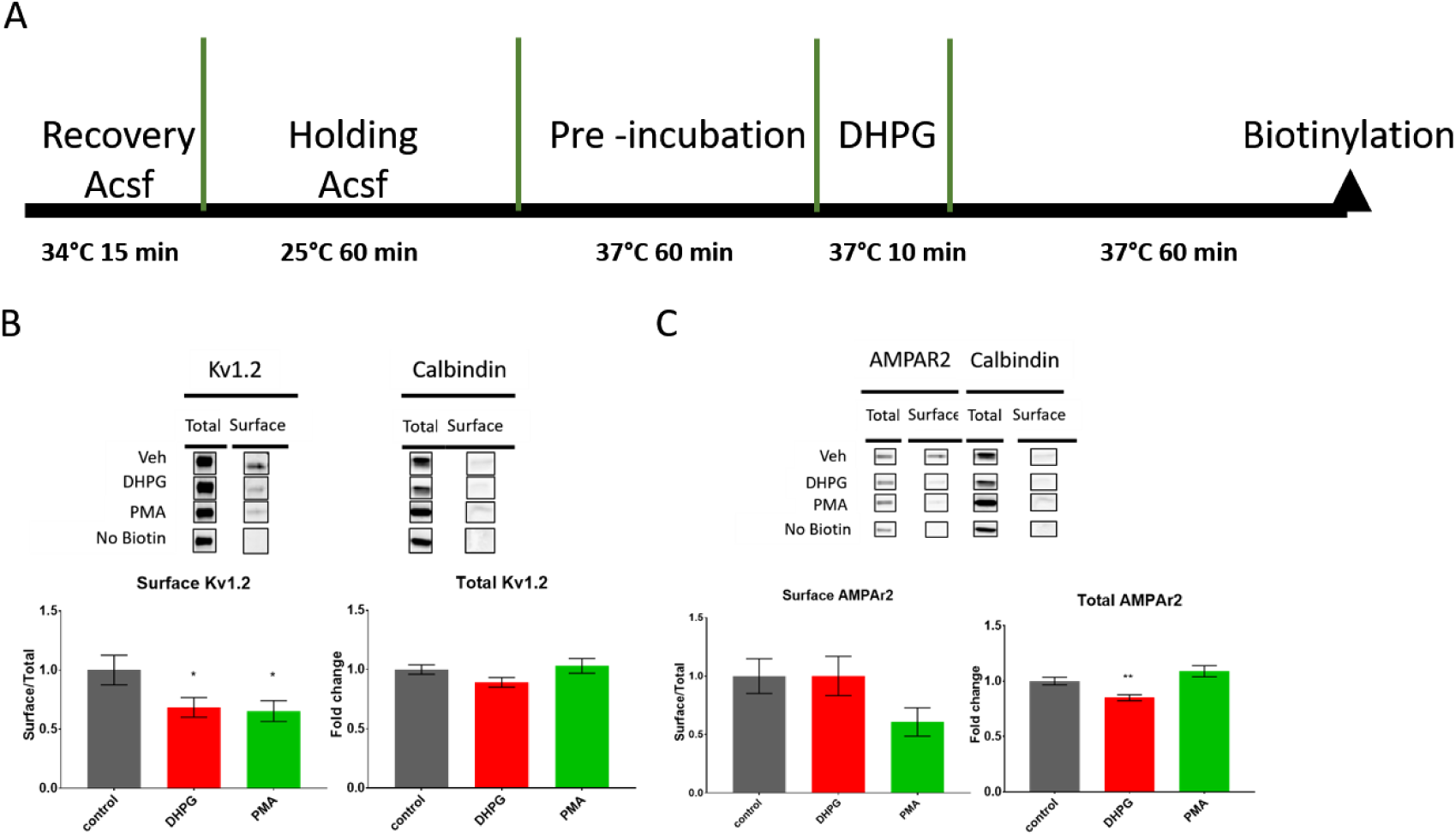
Kv1.2 Endocytosis in the cerebellum. A) A schematic representation of the treatment of Cerebellar slices for biotinylation analysis. B) Cerebellar slices were incubated with 50µM DHPG alone for 10 min or 1µM PMA for 30 min. Then the slices were subjected to biotinylation as mentioned in methods. Western blots (above) and Bar graphs (below) representing changes in the Surface to total ratio of Kv1.2 with various treatments. C) Western blots (above) and Bar graphs (below) representing changes representing changes in the Surface to total ratio of AMPAR2 with various treatments quantified in the same slices as B. * indicates p<0.05 and ** indicates p<0.01 compared to the Control.

**Figure 3:**
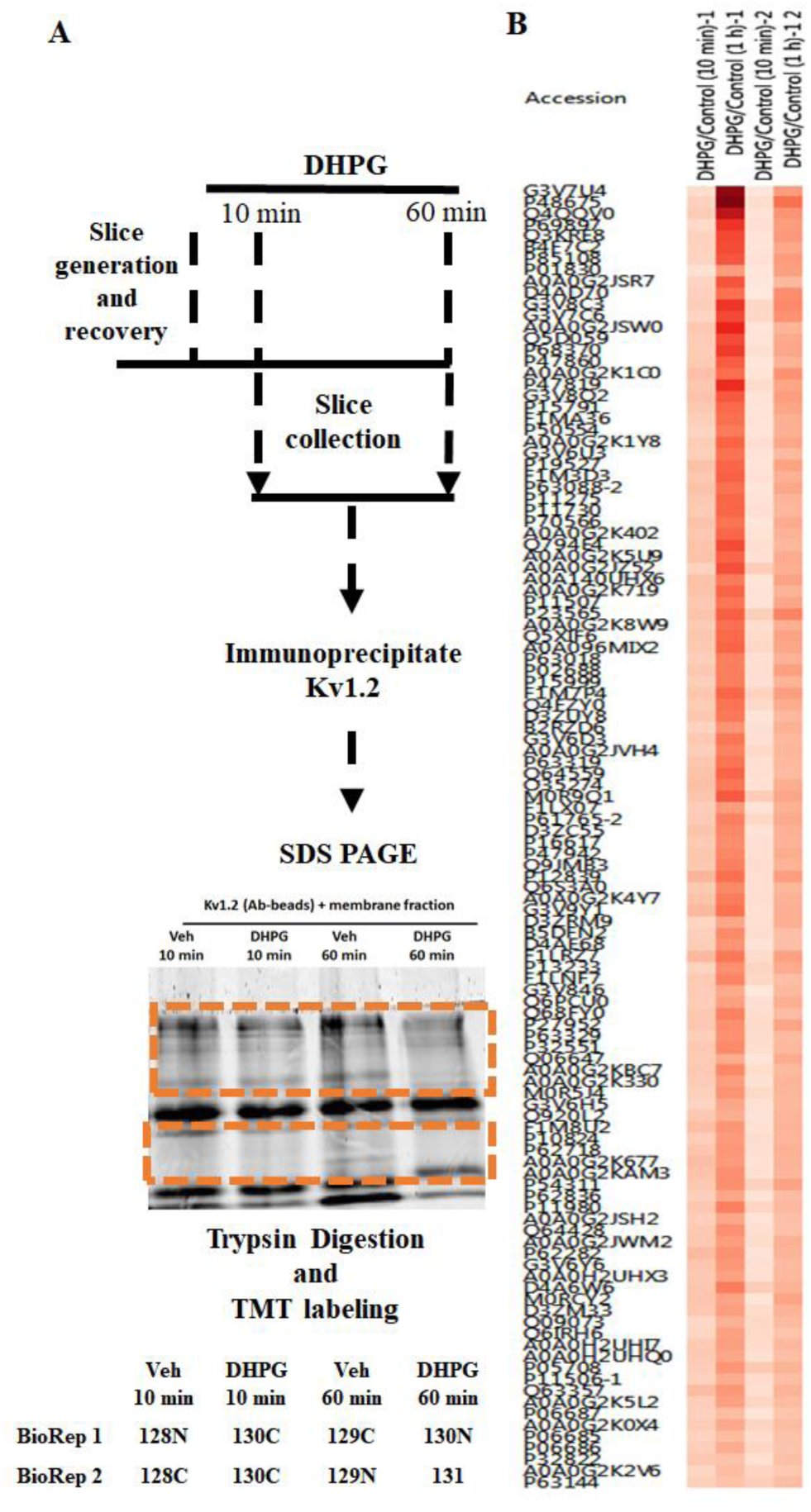
Quantitative Mass spectrometry. A) Schematic representation of the Immunoprecipitation, Gel processing and TMT labels used. The orange boxes show the areas of gel lanes that were processed for TMT labelling, Igg heavy (55KD) and light chain (∼25KD) were excluded from processing. B) Heat map of ratio of proteins (represented by Accession numbers) that Co-ip with Kv1.2 at 10 min and 60 min of DHPG treatment relative to vehicle treated condition. N = 2 biological replicates and 2 technical replicates, pooled from 14 animals.

**Figure 4.**
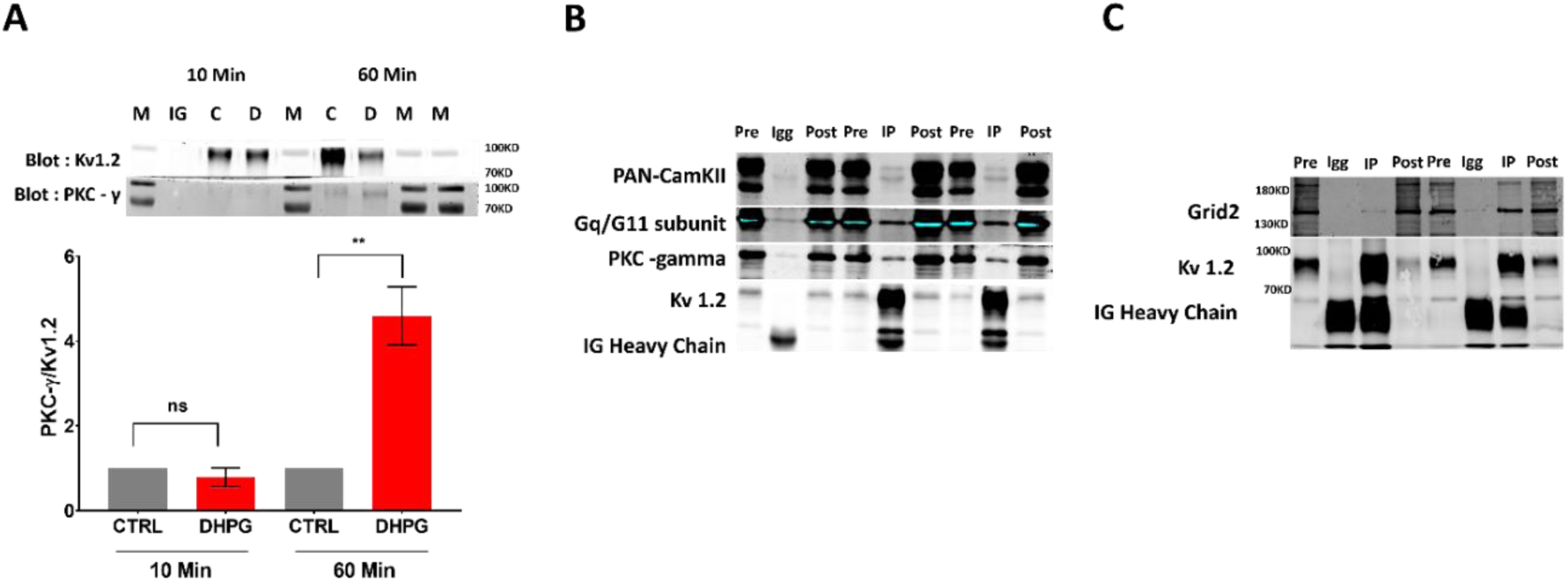
TMT Experiment - western blot validation: A) Western blot of Kv1.2 IP aliquot saved from the TMT mass spectrometry experiment; cerebellar slices treated with DHPG for 10 min or 60 min. The IP was probed for PKC gamma and Kv1.2 (above). The bar graph below shows quantification of PKC gamma relative to Kv1.2. N=2 experiments ** indicates significant difference compared to control. P<0.01 with a one-way ANOVA multiple comparison test. B&C) Independent validation of the Co-ip proteins identified in the MS experiment. Cerebellar lysates were subjected to Kv1.2 immunoprecipitation and were analyzed via western blots with antibodies against CaMKII (PAN-CaMKII antibody), Gq/G11 subunit, PKC-γ, Grid2. Pre – indicates the input, Igg – control Mouse antibody, Post – lysate after immunoprecipitation, IP – immunoprecipitation with anti-Kv1.2 Antibody.

**Figure 5:**
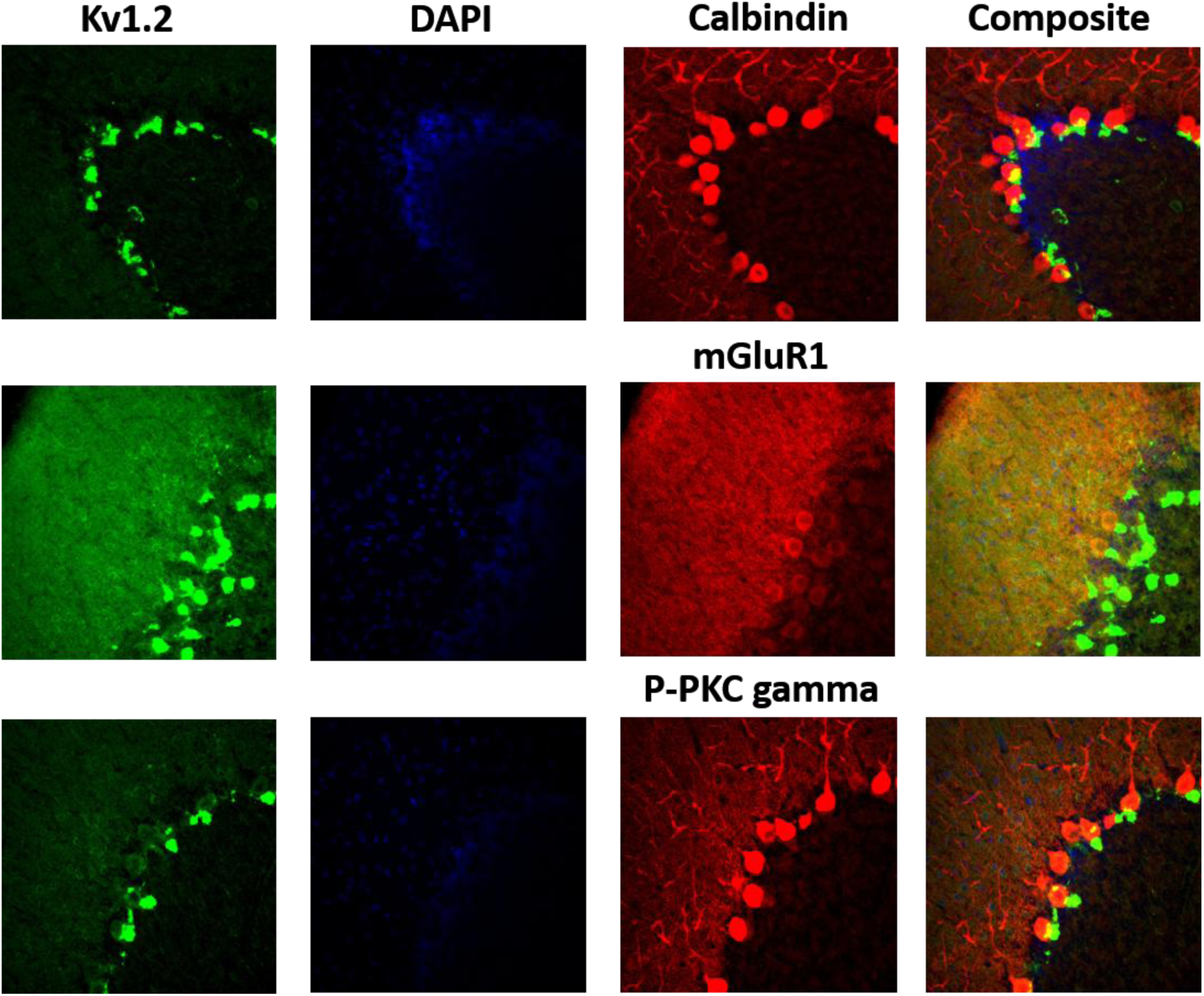
Immunofluorescence images of Rat Cerebellum: Showing Kv1.2, PKC-γ, Calbindin and mGluR1 expression. Cerebellar slice stained with anti-Kv1.2 antibody showing highest expression in the Pinceaux and in molecular layer. Cerebellar slice stained with anti-mGluR1, PKC-γ, Calbindin antibody showing high expression in the Purkinje cell body and in the dendrites in the molecular layer. Composite image showing relative expression of Kv1.2 and mGluR1.

**Figure 6.**
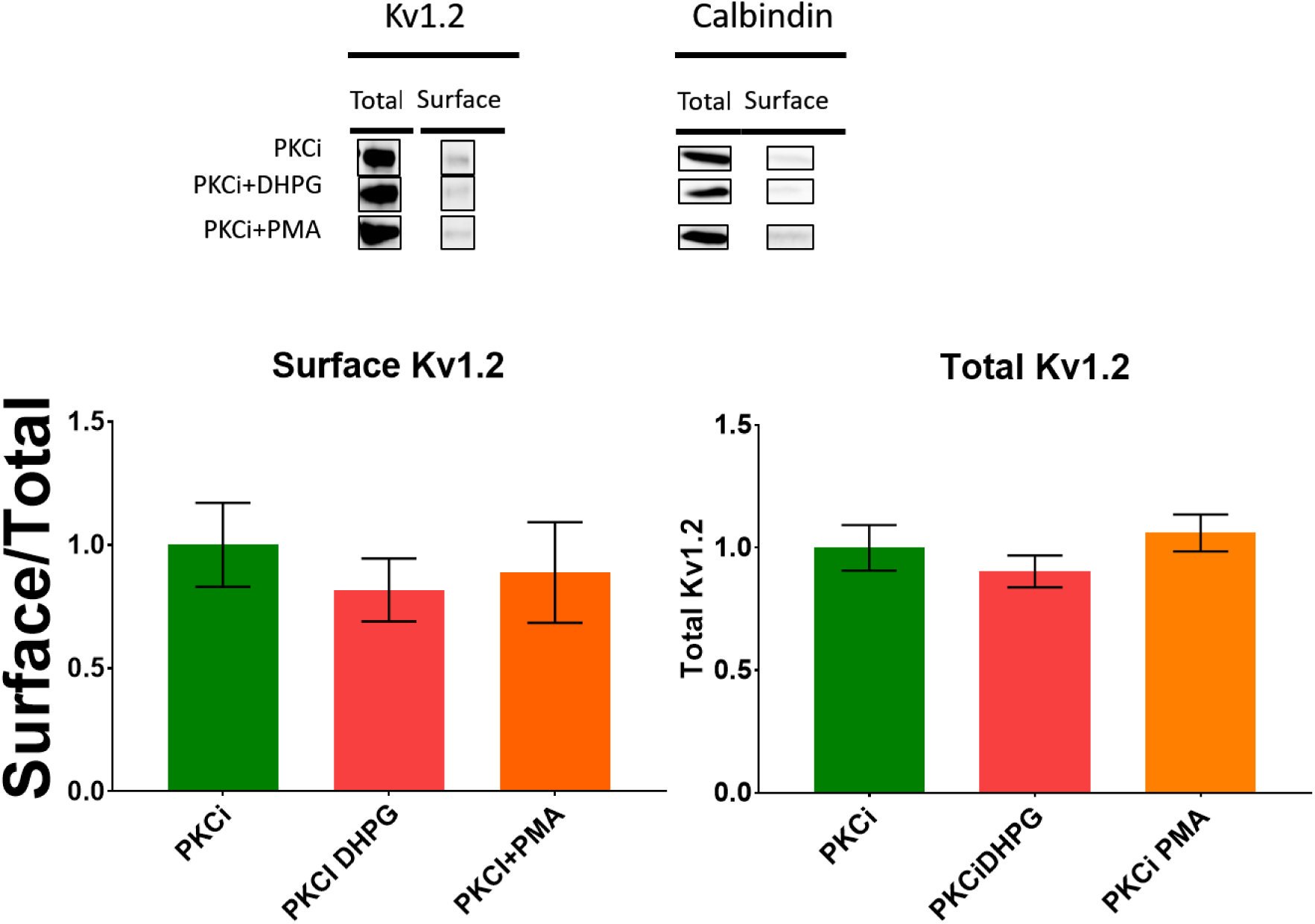
PKC dependent Kv1.2 Endocytosis in the cerebellum. Cerebellar slices were pre-incubated with incubated with PKC inhibitor G06983 for 1 hour prior to treatment with 50µM DHPG alone for 10 min or 1µM PMA for 30 min. Then the slices were subjected to biotinylation as mentioned in methods. Western blots (above) and Bar graphs (below) representing changes in the Surface to total ratio of Kv1.2 with various treatments. Western blots (above) and Bar graphs (below) representing changes representing changes in the Surface to total ratio of Kv1.2.

### mGluR1 activation reduces surface Kv1.2

To more directly test the effect of mGluR1 on Kv1.2, we assessed the effect of the mGluR1 agonist DHPG on Kv1.2 expression at the cell surface. Cerebellar slices were subject to a 10 min application of DHPG (100µM) followed by 1-hour incubation in ACSF (Figure 3A). Surface proteins were labeled using biotinylation, collected onto nutravidin beads and resolved with SDS-PAGE. The level of surface Kv1.2 and AMPA receptor protein was determined by immunoblot. DHPG caused significant reduction in surface Kv1.2 compared to control slices as measured by the ratio intensity of surface to total Kv1.2 signal relative to Vehicle control (DHPG 0.68 ±0.082 vs control; 1 ±0.12; n = 16 slices, N = 4, p = 0.045) (Figure 3B). Because Kv1.2 was previously shown to be regulated by PKC, we used the PKC activator PMA (1µM) as a positive control for Kv1.2 endocytosis. PMA also caused significant loss of surface Kv1.2 (0.65±0.088 vs Control; n=11slices, N=3, p=0.03) (Figure 3B). Total Kv1.2 did not change in slices treated with DHPG or PMA, suggesting that the reduction in surface Kv1.2 is due to effects on channel trafficking. PC depolarization and activation of mGluR1 is known to result in decrease of surface AMPA type ion channel AMPAR2, so we measured the surface levels of AMPAR2 in the same samples. DHPG did not alter surface AMPAR2 levels at 1 hour post stimmulus (Figure 3C) (DHPG 1.001 ±0.168 vs Control; 1 ±0.147; n = 16 slices, N = 4, p = 0.99). In contrast to DHPG, PMA did cause a reduction in surface AMPAR2 channel (0.60±0.12 vs Control; n=11slices, N=3, p=0.0504). Therefore, although mGluR1 can regulate surface expression of both Kv1.2 and AMPAR2 receptors, it can affect each independently.

### Kv1.2 interacts with components of mGluR1 signaling pathway

We next wanted to identify the mechanisms by which mGluR1 regulates Kv1.2. As is true for many proteins, Kv1.2 regulation is in part dictated by its local cellular environment, and in particular by dynamic interaction with proximal regulatory proteins. We therefore used a quantitative mass spectrometry-based proteomics approach to identify components of the cerebellar Kv1.2 protein interaction complex, and to assess mGluR1 mediated changes in those interactions. Cerebellar slices were treated with DHPG (100µM) or vehicle control for 10 min or 1 hour, after which Kv1.2 and its associated protein complex were collected by immunoprecipitation and prepared for analysis by mass spectrometry as described in Methods. Briefly, trypsin digested peptides were labeled with isobaric tandem mass tags, with each tag being specific to an experimental condition. Tagged peptides were then combined prior to mass spectrometry analysis. We have identified 121 proteins that co-immunoprecipitated with Kv1.2 in both biological replicates. Among the proteins identified in the Kv1.2, were those shown in other studies to be in complex with mGluR1 in the cerebellum, including PKC-γ, CaMKII (α, δ isoforms), G_αq_, and GRID2 (20,30).

Interaction with a subset of proteins was increased in slices treated with DHPG for 1h relative to those treated for 10 minutes, PKC-γ, CAMKII (α, δ isoforms), and G_αq_, (fold ratio of (DHPG 1hour/DHPG 10min) >1.5) suggesting an increased binding to Kv1.2 upon mGluR1 activation (Figure 4). In contrast, Kv1.2 interacting proteins likely to be part of the channel itself, inlcuding Kv1.1 and Kv1.6, had fold change ratio < 1.5 with DHPG treatment, suggesting mGluR1 activation does not alter the channel alpha-subunit composition/stoichiometry. Western blot analysis concurred with our MS analysis that the interaction of PKC – γ increases with Kv1.2 in the Post 1-hour DHPG treated samples compared to Vehicle (Figure 4C). We have observed Grid2 Co -ip with Kv1.2 (Figure 4C) very frequently in our mass spectrometry experiments and in one of the biological replicate for TMT experiment, similary we observed PKC-β and PKC-ε in our mass spectromentry experiments infrequently, neither mGluR1, or other mGluR isoforms, were detected in complex with Kv1.2 either by mass spectrometry or by immunoblot.

### Kv1.2, PKC-γ and mGluR1 are expressed in the molecular layer of the cerebellum

Previous studies have shown that mGluR1 is expressed in the molecular layer of cerebellum (20,31–34). Anton N. Shuvaev *et al* (Figure 2D in Reference (33)) show that mGluR1 is primarily localized in PCs in the molecular layer and Armburst *et al* performed cluster analysis of mGluR1 and Calbindin in mouse cerebellum and show that mGluR1 is localized in the Purkinje cell dendrites (32) Figure 5). Separate studies showed that Kv1.2 is expressed in the molecular layer of the cerebellar cortex both in the PC dendrites and BC axon terminals (1,2,35,36). Here, we used immunofluorescence to determine the areas of overlap between Kv1.2, mGluR1 and the mGluR1 signaling protein PKC-γ in parasagittal sections from adult rat cerebellum. Consistent with our previous observations (13) and with multiple earlier studies(1), Kv1.2 expression was detected in the molecular layer and in the BC axon terminals (Figure 6). In agreement with previous studies we observed that the relative abundance of mGluR1 appears higher in the molecular layer than in the BC axon terminal (20,34). We also observe that similar to mGluR1, PKC-γ is highly expressed in the PCs, but not in BC axon terminals (Figure 6). We therefore conclude that modulation of Kv1.2 originates from mGluR1 expressed in PCs and most likely involves Kv1.2 expressed in PCs as well.

### Broad spectrum PKC inhibitor attenuates mGluR1 mediated loss of surface Kv1.2

Since mGluR1 activation causes endocytosis of ion channels via PKC, we used a broad-spectrum PKC inhibitor Go6983 which blocks all calcium dependent PKC but not a typical PKC isoforms. In the presence of the PKC inhibitor (PKCi in Figures) neither DHPG nor PMA caused further loss of surface Kv1.2 (Figure 7) (PKCi DHPG 0.81 ± 0.1276, PKCi PMA 0.88±0.2038 vs PKCi 1.0 ± 0.17, p = 0.404, n= 8 slices, N = 2). We conclude that mGluR1 modulation of surface Kv1.2 proceeds in part through a PKC dependent mechanism, possibly involving PKC-γ.

## Discussion

One of the most well-known mechanisms for learning-related synaptic plasticity in the cerebellum involves cerebellar LTD. During cerebellar LTD, activation of mGluR1 occurs, which through its canonical signaling pathway activates PKC, which results in phosphorylation and endocytosis of AMPAR2 thus resulting in LTD. Knock out of mGluR1 impairs EBC, a cerebellar form of learning and memory, and mGluR1 rescue in PCs restores EBC (18,19). Schonewille *et al*. showed that in mutant mice with impaired AMPAR2 endocytosis and consequently impaired parallel fiber-PC LTD, EBC is normal, suggesting that AMPAR trafficking is involved in but not required for EBC. Further, Yamaguchi *et al*. were able to induce LTD in cerebellar slices in AMPAR2 endocytosis deficient mutants via modified stimulation protocols, and that PKCα is required for this LTD, again suggesting that mGluR1 signaling is essential but might involve mechanisms beyond AMPAR2 endocytosis (37,38). Overall, this suggests that for normal EBC to occur, mGluR1 activation is required, but AMPAR2 endocytosis is not essential and there may be other mechanisms that might be involved.

We have previously shown that Kv1.2 is involved in cerebellar learning and memory. Pharmacological inhibition of Kv1.2 facilitates EBC and EBC itself modulates surface Kv1.2 expression (13,14,23). We have shown that direct activation of mGluR1 by infusion of DHPG into the cerebellum improves EBC acquisition. Here we show that application of glutamate and high potassium to cerebellar slices results in increase in tyrosine phosphorylation of proteins immunoprecipitated with an anti-Kv1.2 antibody, and an mGluR1 antagonist prevented that increase in tyrosine phosphorylation. Tyrosine phosphorylation of Kv1.2 is associated with channel endocytosis (39) and a resultant decrease in channel function. It is therefore consistent with the finding that DHPG reduces surface Kv1.2 levels in cerebellar slices. Since pharmacological inhibition of Kv1.2 enhances EBC (13), such a reduction in surface Kv1.2 is also consistent with our finding that mGluR1 activation with DHPG enhances EBC. However, Kv1.2 is likely to be regulated by phosphorylation of multiple tyrosine’s (9,12) and their roles in Kv1.2 regulation in specific cerebellar neurons remains to be determined. Further, Kv1.2 surface expression and function are regulated by serine/threonine phosphorylation as well (11,28). Nevertheless, the findings presented here are consistent with model in which mGluR1 modulates Kv1.2 function in part by affecting the channel’s post-translational modification.

One of the consequences of Kv1.2 post-translational modification is modulation of its interaction with other proteins. For example, in HEK293 cells and in GST-pull down studies, tyrosine phosphorylation of Kv1.2 decreases its interaction with the actin regulating protein cortactin (40). This in turn promotes endocytosis of the channel from the cell surface.

In this study we therefore sought to identify protein interactors of Kv1.2 as a way to shed light on the potential pathways that might be involved in its regulation by mGluR1. In the TMT mass spectrometry study summarized in supplementary table, several proteins are particularly significant because they are known to also interact with mGluR1(41). These include PKC – γ and calmodulin kinases (multiple isoforms) were shown physically interact with Kv1.2 (41). We note that we did not find mGluR1 in complex with immunoprecipitated Kv1.2 in either immunoblot or mass spectrometry analyses. Whether this is because mGluR1 does not interact directly with Kv1.2, or because the interaction is too unstable to detect under the immunoprecipitation conditions used in this study remains to be determined.

The DHPG induced reduction of surface Kv1.2 levels in cerebellar slices was mimicked by the PKC activator PMA and reduced by the PKC inhibitor Go6983. This suggests that the canonical mGluR1 – PKC signaling mechanism might be involved in the regulation of Kv1.2. Whether this involves PKC-γ or another PKC isoform not detected in our mass spectrometry analysis and remains to be determined. It is possible, for example, that the multiple isoforms of PKC that are activated by mGluR1 might have different targets, with PKC-gamma targeting Kv1.2 and PKC-alpha targeting AMPAR2 (42–44). Hirono *et al* have shown that PKCα and PKCβI isoforms translocate to the PC dendrites upon stimulation with high K^+^ - Glutamate stimulus while PKC γ did not. Leitges *et al* have shown that LTD is absent in PKC α deficient cerebellum and was rescued only with the expression of the α isoform indicating that α isoform of PKC is more important for LTD and there for AMPAR2 endocytosis (42,43). Such a model would be consistent with our finding that in our studies DHPG did not affect AMPAR2 surface levels while global activation of diacylglycerol sensitive PKC isoforms with PMA did (44).

mGluR1 and PKC-γ are mostly expressed in the molecular layer within the PC cell body and dendrites. Little evidence exists for their expression on basket cell soma or axon terminals. Consistent with this, we detected expression of mGluR1 and PKC-gamma in the molecular layer but not in pinceau’s (Figure 6). It is therefore likely that mGluR1 modulates Kv1.2 in PC dendrites and not in the BC axon terminals. However, it is possible that the regulation of Kv1.2 by mGluR1 is indirect. We have previously shown that secretin receptors regulate Kv1.2, secretin is released from the PCs upon depolarization(45), therefore it is possible that mGluR1 activation which results in increased Ca levels induces secretin release from PCs and the released secretin via secretin receptors on PCs and basket cell axons down regulate Kv1.2. It is essential therefore to study the activation of mGluR1 when secretin receptors are inhibited on Kv1.2 levels and on AMPAR2. Regardless of the mechanisms by which mGluR1 modulates Kv1.2, Suppression of Kv1.2 channels in the PC could lead to increased excitability and thereby facilitate LTD (41) by increasing calcium influx. While endocytosis of Kv1.2 at the BC results in increased GABAergic input on the AIS of PC there by suppressing PC (3,13). The net effect of enhanced LTD and/or increased GABAergic inhibition would disinhibit the deep cerebellar nuclei and might help improve EBC.

## Acknowledgements

This work is done by UVM internal Grant to AM, JG. The proteomics work was supported in part by Vermont Genetics Network Proteomics Facility is supported through NIH grant P20GM103449 from the INBRE Program of the National Institute of General Medical Sciences.

Dr Ying Wai Lam contributed to design, execution and data analysis for TMT mass spectrometry experiments as Part of proteomics facility, UVM.

## Author Contributions

SM, AD Contributed to original ideas, SM, AD contributed to design, execution and data analysis for Figures 1-6. SM,

## Conflicts of Interest

None

## Notes

### Competing Interest Statement

The authors have declared no competing interest.

### Summary of Updates

EBC Behavioral experiments were published separately and to EBC mentions as part of this manuscript are removed in this version; Few grammatical errors were corrected; Figure numbers were formatted in results section.

## References

1. Kole, M. J., Qian, J., Waase, M. P., Klassen, T. L., Chen, T. T., Augustine, G. J., and Noebels, J. L. (2015) Selective Loss of Presynaptic Potassium Channel Clusters at the Cerebellar Basket Cell Terminal Pinceau in *Adam11* Mutants Reveals Their Role in Ephaptic Control of Purkinje Cell Firing. The Journal of Neuroscience 35, 11433–11444

2. Williams, M. R., Fuchs, J. R., Green, J. T., and Morielli, A. D. (2012) Cellular mechanisms and behavioral consequences of Kv1.2 regulation in the rat cerebellum. The Journal of Neuroscience 32, 9228–9237

3. Xie, G., Harrison, J., Clapcote, S. J., Huang, Y., Zhang, J. Y., Wang, L. Y., and Roder, J. C. (2010) A new Kv1.2 channelopathy underlying cerebellar ataxia. The Journal of biological chemistry 285, 32160–32173

4. Brew, H. M., Gittelman, J. X., Silverstein, R. S., Hanks, T. D., Demas, V. P., Robinson, L. C., Robbins, C. A., McKee-Johnson, J., Chiu, S. Y., Messing, A., and Tempel, B. L. (2007) Seizures and reduced life span in mice lacking the potassium channel subunit Kv1.2, but hypoexcitability and enlarged Kv1 currents in auditory neurons. Journal of neurophysiology 98, 1501–1525

5. Robbins, C. A., and Tempel, B. L. (2012) Kv1.1 and Kv1.2: Similar channels, different seizure models. Epilepsia 53, 134–141

6. Kearney, J. A. (2015) KCNA2-Related Epileptic Encephalopathy. Pediatric neurology briefs 29, 27

7. Pena, S. D., and Coimbra, R. L. (2015) Ataxia and myoclonic epilepsy due to a heterozygous new mutation in KCNA2: proposal for a new channelopathy. Clinical genetics 87, e1–3

8. Allou, L., Julia, S., Amsallem, D., El Chehadeh, S., Lambert, L., Thevenon, J., Duffourd, Y., Saunier, A., Bouquet, P., Pere, S., Moustaine, A., Ruaud, L., Roth, V., Jonveaux, P., and Philippe, C. (2016) Rett-like phenotypes: expanding the genetic heterogeneity to the KCNA2 gene and first familial case of CDKL5-related disease. Clinical genetics

9. Huang, X. Y., Morielli, A. D., and Peralta, E. G. (1993) Tyrosine kinase-dependent suppression of a potassium channel by the G protein-coupled m1 muscarinic acetylcholine receptor. Cell 75, 1145–1156

10. Hattan, D., Nesti, E., Cachero, T. G., and Morielli, A. D. (2002) Tyrosine phosphorylation of Kv1.2 modulates its interaction with the actin-binding protein cortactin. The Journal of biological chemistry 277, 38596–38606

11. Connors, E. C., Ballif, B. A., and Morielli, A. D. (2008) Homeostatic regulation of Kv1.2 potassium channel trafficking by cyclic AMP. The Journal of biological chemistry 283, 3445–3453

12. Nesti, E., Everill, B., and Morielli, A. D. (2004) Endocytosis as a mechanism for tyrosine kinase-dependent suppression of a voltage-gated potassium channel. Molecular biology of the cell 15, 4073–4088

13. Williams, M. R., Fuchs, J. R., Green, J. T., and Morielli, A. D. (2012) Cellular mechanisms and behavioral consequences of Kv1.2 regulation in the rat cerebellum. The Journal of neuroscience : the official journal of the Society for Neuroscience 32, 9228–9237

14. Fuchs, J. R., Darlington, S. W., Green, J. T., and Morielli, A. D. (2017) Cerebellar learning modulates surface expression of a voltage-gated ion channel in cerebellar cortex. Neurobiology of learning and memory 142, 252–262

15. Nakao, H., Nakao, K., Kano, M., and Aiba, A. (2007) Metabotropic glutamate receptor subtype-1 is essential for motor coordination in the adult cerebellum. Neuroscience research 57, 538–543

16. Ichise, T., Kano, M., Hashimoto, K., Yanagihara, D., Nakao, K., Shigemoto, R., Katsuki, M., and Aiba, A. (2000) mGluR1 in cerebellar Purkinje cells essential for long-term depression, synapse elimination, and motor coordination. Science (New York, N.Y.) 288, 1832–1835

17. Aiba, A., Kano, M., Chen, C., Stanton, M. E., Fox, G. D., Herrup, K., Zwingman, T. A., and Tonegawa, S. (1994) Deficient cerebellar long-term depression and impaired motor learning in mGluR1 mutant mice. Cell 79, 377–388

18. Kishimoto, Y., Fujimichi, R., Araishi, K., Kawahara, S., Kano, M., Aiba, A., and Kirino, Y. (2002) mGluR1 in cerebellar Purkinje cells is required for normal association of temporally contiguous stimuli in classical conditioning. The European journal of neuroscience 16, 2416–2424

19. Ohtani, Y., Miyata, M., Hashimoto, K., Tabata, T., Kishimoto, Y., Fukaya, M., Kase, D., Kassai, H., Nakao, K., Hirata, T., Watanabe, M., Kano, M., and Aiba, A. (2014) The synaptic targeting of mGluR1 by its carboxyl-terminal domain is crucial for cerebellar function. The Journal of neuroscience : the official journal of the Society for Neuroscience 34, 2702–2712

20. Kato, A. S., Knierman, M. D., Siuda, E. R., Isaac, J. T. R., Nisenbaum, E. S., and Bredt, D. S. (2012) Glutamate Receptor δ2 Associates with Metabotropic Glutamate Receptor 1 (mGluR1), Protein Kinase Cγ, and Canonical Transient Receptor Potential 3 and Regulates mGluR1-Mediated Synaptic Transmission in Cerebellar Purkinje Neurons. The Journal of Neuroscience 32, 15296–15308

21. Ting, J. T., Daigle, T. L., Chen, Q., and Feng, G. (2014) Acute brain slice methods for adult and aging animals: application of targeted patch clamp analysis and optogenetics. Methods in molecular biology (Clifton, N.J.) 1183, 221–242

22. Titley, H. K., Heskin-Sweezie, R., and Broussard, D. M. (2010) The Bidirectionality of Motor Learning in the Vestibulo-ocular Reflex Is a Function of Cerebellar mGluR1 Receptors. 104, 3657–3666

23. Fuchs, J. R., Robinson, G. M., Dean, A. M., Schoenberg, H. E., Williams, M. R., Morielli, A. D., and Green, J. T. (2014) Cerebellar secretin modulates eyeblink classical conditioning. Learning & memory (Cold Spring Harbor, N.Y.) 21, 668–675

24. Tanaka, K., and Augustine, G. J. (2008) A positive feedback signal transduction loop determines timing of cerebellar long-term depression. Neuron 59, 608–620

25. Chae, H. G., Ahn, S. J., Hong, Y. H., Chang, W. S., Kim, J., and Kim, S. J. (2012) Transient Receptor Potential Canonical Channels Regulate the Induction of Cerebellar Long-Term Depression. The Journal of Neuroscience 32, 12909–12914

26. Erkens, M., Tanaka-Yamamoto, K., Cheron, G., Marquez-Ruiz, J., Prigogine, C., Schepens, J. T., Nadif Kasri, N., Augustine, G. J., and Hendriks, W. J. (2015) Protein tyrosine phosphatase receptor type R is required for Purkinje cell responsiveness in cerebellar long-term depression. Molecular brain 8, 1

27. Kohda, K., Kakegawa, W., Matsuda, S., Yamamoto, T., Hirano, H., and Yuzaki, M. (2013) The delta2 glutamate receptor gates long-term depression by coordinating interactions between two AMPA receptor phosphorylation sites. Proceedings of the National Academy of Sciences of the United States of America 110, E948–957

28. Yang, J. W., Vacher, H., Park, K. S., Clark, E., and Trimmer, J. S. (2007) Trafficking-dependent phosphorylation of Kv1.2 regulates voltage-gated potassium channel cell surface expression. Proceedings of the National Academy of Sciences of the United States of America 104, 20055–20060

29. Stirling, L., Williams, M. R., and Morielli, A. D. (2009) Dual roles for RHOA/RHO-kinase in the regulated trafficking of a voltage-sensitive potassium channel. Molecular biology of the cell 20, 2991–3002

30. Jin, D.-Z., Guo, M.-L., Xue, B., Fibuch, E. E., Choe, E. S., Mao, L.-M., and Wang, J. Q. (2013) Phosphorylation and Feedback Regulation of Metabotropic Glutamate Receptor 1 by CaMKII. The Journal of neuroscience : the official journal of the Society for Neuroscience 33, 3402–3412

31. Mitsumura, K., Hosoi, N., Furuya, N., and Hirai, H. (2011) Disruption of metabotropic glutamate receptor signalling is a major defect at cerebellar parallel fibre-Purkinje cell synapses in staggerer mutant mice. J Physiol 589, 3191–3209

32. Armbrust, K. R., Wang, X., Hathorn, T. J., Cramer, S. W., Chen, G., Zu, T., Kangas, T., Zink, A. N., Öz, G., Ebner, T. J., and Ranum, L. P. W. (2014) Mutant β-III Spectrin Causes mGluR1α Mislocalization and Functional Deficits in a Mouse Model of Spinocerebellar Ataxia Type 5. The Journal of Neuroscience 34, 9891–9904

33. Shuvaev, A. N., Hosoi, N., Sato, Y., Yanagihara, D., and Hirai, H. (2017) Progressive impairment of cerebellar mGluR signalling and its therapeutic potential for cerebellar ataxia in spinocerebellar ataxia type 1 model mice. The Journal of Physiology 595, 141–164

34. Kurnellas, M. P., Lee, A. K., Szczepanowski, K., and Elkabes, S. (2007) Role of plasma membrane calcium ATPase isoform 2 in neuronal function in the cerebellum and spinal cord. Annals of the New York Academy of Sciences 1099, 287–291

35. Iwakura, A., Uchigashima, M., Miyazaki, T., Yamasaki, M., and Watanabe, M. (2012) Lack of Molecular–Anatomical Evidence for GABAergic Influence on Axon Initial Segment of Cerebellar Purkinje Cells by the Pinceau Formation. The Journal of Neuroscience 32, 9438–9448

36. Buttermore, E. D., Thaxton, C. L., and Bhat, M. A. (2013) Organization and maintenance of molecular domains in myelinated axons. Journal of Neuroscience Research 91, 603–622

37. Schonewille, M., Gao, Z., Boele, H. J., Veloz, M. F., Amerika, W. E., Simek, A. A., De Jeu, M. T., Steinberg, J. P., Takamiya, K., Hoebeek, F. E., Linden, D. J., Huganir, R. L., and De Zeeuw, C. I. (2011) Reevaluating the role of LTD in cerebellar motor learning. Neuron 70, 43–50

38. Yamaguchi, K., Itohara, S., and Ito, M. (2016) Reassessment of long-term depression in cerebellar Purkinje cells in mice carrying mutated GluA2 C terminus. Proceedings of the National Academy of Sciences of the United States of America 113, 10192–10197

39. Hyun, J. H., Eom, K., Lee, K. H., Ho, W. K., and Lee, S. H. (2013) Activity-dependent downregulation of D-type K+ channel subunit Kv1.2 in rat hippocampal CA3 pyramidal neurons. The Journal of physiology 591, 5525–5540

40. Williams, M. R., Markey, J. C., Doczi, M. A., and Morielli, A. D. (2007) An essential role for cortactin in the modulation of the potassium channel Kv1.2. Proceedings of the National Academy of Sciences of the United States of America 104, 17412–17417

41. Kato, A. S., Knierman, M. D., Siuda, E. R., Isaac, J. T., Nisenbaum, E. S., and Bredt, D. S. (2012) Glutamate receptor delta2 associates with metabotropic glutamate receptor 1 (mGluR1), protein kinase Cgamma, and canonical transient receptor potential 3 and regulates mGluR1-mediated synaptic transmission in cerebellar Purkinje neurons. The Journal of neuroscience : the official journal of the Society for Neuroscience 32, 15296–15308

42. Leitges, M., Kovac, J., Plomann, M., and Linden, D. J. (2004) A unique PDZ ligand in PKCalpha confers induction of cerebellar long-term synaptic depression. Neuron 44, 585–594

43. Hirono, M., Sugiyama, T., Kishimoto, Y., Sakai, I., Miyazawa, T., Kishio, M., Inoue, H., Nakao, K., Ikeda, M., Kawahara, S., Kirino, Y., Katsuki, M., Horie, H., Ishikawa, Y., and Yoshioka, T. (2001) Phospholipase Cbeta4 and protein kinase Calpha and/or protein kinase CbetaI are involved in the induction of long term depression in cerebellar Purkinje cells. The Journal of biological chemistry 276, 45236–45242

44. Linden, D. J., and Connor, J. A. (1991) Participation of postsynaptic PKC in cerebellar long-term depression in culture. *Science (New York*, N.Y.) 254, 1656–1659

45. Yung, W.-H., Leung, P.-S., Ng, S. S. M., Zhang, J., Chan, S. C. Y., and Chow, B. K. C. (2001) Secretin Facilitates GABA Transmission in the Cerebellum. 21, 7063–7068

